# Streamlining Data-Intensive Biology With Workflow Systems

**DOI:** 10.1101/2020.06.30.178673

**Authors:** Taylor Reiter, Phillip T. Brooks, Luiz Irber, Shannon E.K. Joslin, Charles M. Reid, Camille Scott, C. Titus Brown, N. Tessa Pierce

## Abstract

As the scale of biological data generation has increased, the bottleneck of research has shifted from data generation to analysis. Researchers commonly need to build computational workflows that include multiple analytic tools and require incremental development as experimental insights demand tool and parameter modifications. These workflows can produce hundreds to thousands of intermediate files and results that must be integrated for biological insight. Data-centric workflow systems that internally manage computational resources, software, and conditional execution of analysis steps are reshaping the landscape of biological data analysis, and empowering researchers to conduct reproducible analyses at scale. Adoption of these tools can facilitate and expedite robust data analysis, but knowledge of these techniques is still lacking. Here, we provide a series of practices and strategies for leveraging workflow systems with structured project, data, and resource management to streamline large-scale biological analysis. We present these strategies in the context of high-throughput sequencing data analysis, but the principles are broadly applicable to biologists working beyond this field.

**Author Summary:** We present a guide for workflow-enabled biological sequence data analysis, developed through our own teaching, training and analysis projects. We recognize that this is based on our own use cases and experiences, but we hope that our guide will contribute to a larger discussion within the open source and open science communities and lead to more comprehensive resources. Our main goal is to accelerate the research of scientists conducting sequence analyses by introducing them to organized workflow practices that not only benefit their own research but also facilitate open and reproducible science.

## Introduction

Biological research has become increasingly computational. In particular, genomics has experienced a deluge of high-throughput sequencing data that has already reshaped our understanding of the diversity and function of organisms and communities, building basic understanding from ecosystems to human health. The analysis workflows used to produce these insights often integrate hundreds of steps and involve a myriad of decisions ranging from small-scale tool and parameter choices to larger-scale design decisions around data processing and statistical analyses. Each step relies not just on analysis code written by the researcher, but on third-party software, its dependencies, and the compute infrastructure and operating system on which the code is executed. Historically, this has led to the patchwork availability of underlying code for analyses as well as a lack of interoperability of the resulting software and analysis pipelines across compute systems [1]. Combined with unmet training needs in biological data analysis, these conditions undermine the reuse of data and the reproducibility of biological research, vastly limiting the value of our generated data [2].

The biological research community is strongly committed to addressing these issues, recently formalizing the FAIR practices: the idea that all life sciences research (including data and analysis workflows) should be Findable, Accessible, Interoperable, and Reusable [3]. For computational analyses, these ideals are readily achievable with current technologies, but implementing them in practice has proven difficult, particularly for biologists with little training in computing [3]. However, the recent maturation of data-centric workflow systems designed to automate and facilitate computational workflows is expanding our capacity to conduct end-to-end FAIR analyses [5]. These workflow systems are designed to handle some aspects of computational workflows internally: namely, the interactions with software and computing infrastructure, and the ordered execution of each step of an analysis. By reducing the manual input and monitoring required at each analysis juncture, these integrated systems ensure that analyses are repeatable and can be executed at much larger scales. In concert, the standardized information and syntax required for rule-based workflow specification makes code inherently modular and more easily transferable between projects [5,6]. For these reasons, workflow systems are rapidly becoming the workhorses of modern bioinformatics.

Adopting workflow systems requires some level of up-front investment, first to understand the structure of the system, and then to learn the workflow-specific syntax. These challenges can preclude adoption, particularly for researchers without significant computational experience [4]. In our experiences with both research and training, these initial learning costs are similar to those required for learning more traditional analysis strategies, but then provide a myriad of additional benefits that both facilitate and accelerate research. Furthermore, online communities for sharing reusable workflow code have proliferated, meaning the initial cost of encoding a workflow in a system is mitigated via use and re-use of common steps, leading to faster time-to-insight [5,7].

Building upon the rich literature of “best” and “good enough” practices for computational biology [8,9,10], we present a series of strategies and practices for adopting workflow systems to streamline data-intensive biology research. This manuscript is designed to help guide biologists towards project, data, and resource management strategies that facilitate and expedite reproducible data analysis in their research. We present these strategies in the context of our own experiences working with high-throughput sequencing data, but many are broadly applicable to biologists working beyond this field.

## Worklows facilitate data-intensive biology

Data-intensive biology typically requires that researchers execute computational workflows using multiple analytic tools and apply them to many experimental samples in a systematic manner. These workflows commonly produce hundreds to thousands of intermediate files and require incremental changes as experimental insights demand tool and parameter modifications. Many intermediate steps are central to the biological analysis, but others, such as converting between file formats, are rote computational tasks required to passage data from one tool to the next. Some of these steps can fail silently, producing incomplete intermediate files that imperceptively invalidate downstream results and biological inferences. Properly managing and executing all of these steps is vital, but can be both time-consuming and error-prone, even when automated with scripting languages such as bash.

The emergence and maturation of workflow systems designed with bioinformatic challenges in mind has revolutionized computing in data intensive biology [11]. Workflow systems contain powerful infrastructure for workflow management that can coordinate runtime behavior, self-monitor progress and resource usage, and compile reports documenting the results of a workflow (**Figure 1**). These features ensure that the steps for data analysis are repeatable and at least minimally described from start to finish. When paired with proper software management, fully-contained workflows are scalable, robust to software updates, and executable across platforms, meaning they will likely still execute the same set of commands with little investment by the user after weeks, months, or years.

**Figure 1:**
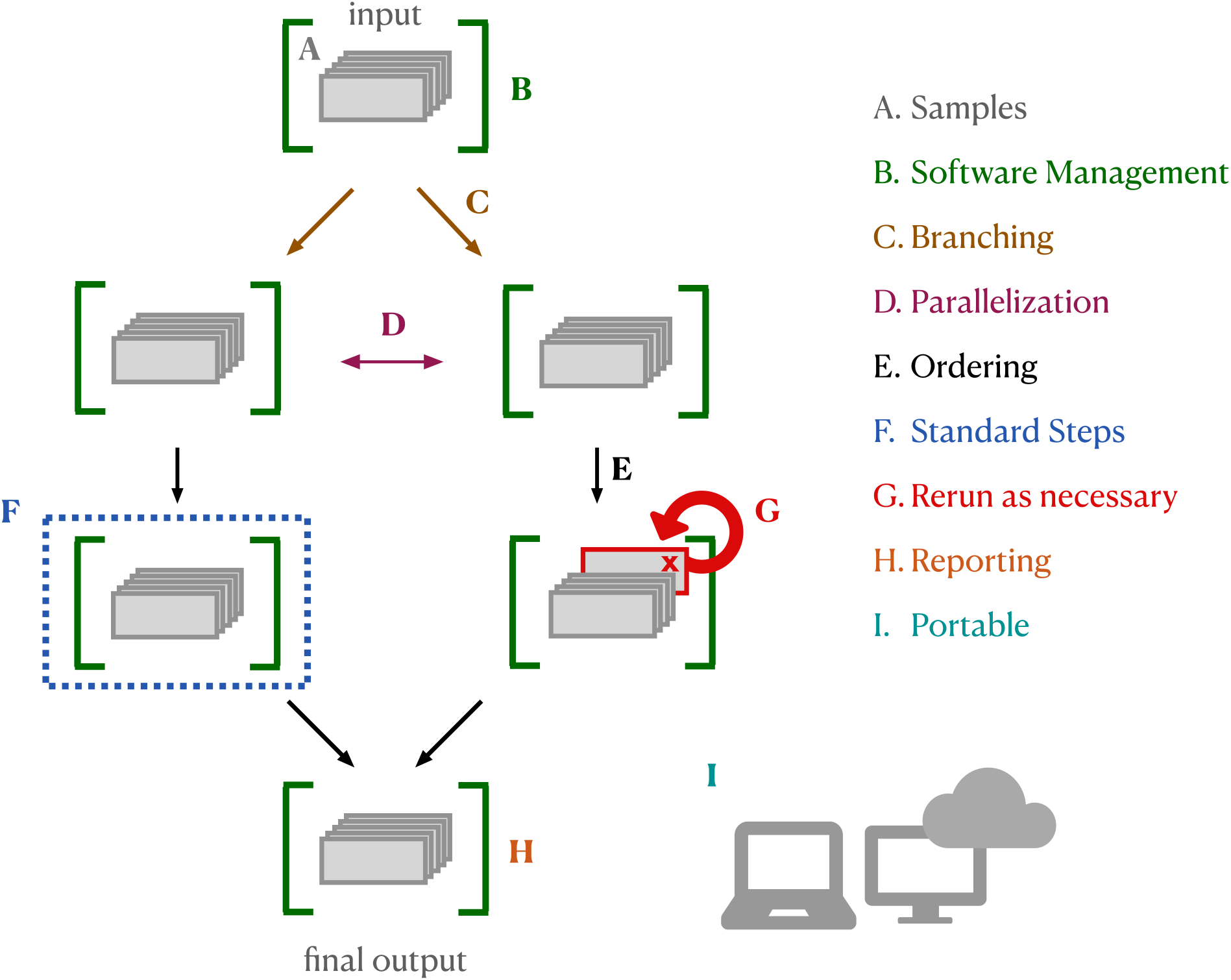
Worklow Systems: Bioinformatic workflow systems have built-in functionality that facilitates and simplifies running analysis pipelines. **A. Samples:** Workflow systems enable you to use the same code to run each step on each sample. Samples can be easily added if the analysis expands. **B. Software Management:** Integration with software management tools (e.g. conda, singularity, docker) can automate software installation for each step. **C. Branching, D. Parallelization, and E. Ordering:** Workflow systems handle conditional execution, ensuring that tasks are executed in the correct order for each sample file, including executing independent steps in parallel if possible given the resources provided. **F. Standard Steps:** Many steps are now considered “standard” (e.g. quality control). Workflow languages keep all information for a step together and can be written to enable you to remix and reuse individual steps across pipelines. **G. Rerun as necessary:** Workflow systems keep track of which steps executed properly and on which samples, and allow you to rerun failed steps (or additional steps) rather than re-executing the entire workflow. **H. Reporting:** Workflow languages enable comprehensive reporting on workflow execution and resource utilization by each tool. **I. Portability:** Analyses written in workflow languages (with integrated software management) can be run across computing systems without changes to code.

To properly direct an analysis, workflow systems need to encode information about the relationships between every workflow step. In practice, this means that each analysis step must specify the input (or types of inputs) needed for that step, and the output (or types of outputs) being produced. This structure provides several additional benefits. First, workflows become minimally self-documented, as the directed graph produced by workflow systems can be exported and visualized, producing a graphical representation of the relationships between all steps in a pipeline (see **Figure 5**). Next, workflows are more likely to be fully enclosed without undocumented steps that are executed by hand, meaning analyses are more likely to be reproducible. Finally, each step becomes a self-contained unit that can be used and re-used across multiple analysis workflows, so scientists can spend less time implementing standard steps, and more time on their specific research questions. In sum, the internal scaffolding provided by workflow systems helps build analyses that are generally better documented, repeatable, transferable, and scalable.

### Getting started with worklows

The workflow system you choose will be largely dependent on your analysis needs. Here, we draw a distinction between two types of workflows: “research” workflows that are under iterative development to answer novel scientific questions, and “production” workflows, which have reached maturity and are primarily used to run a standard analysis on new samples. In particular, research workflows require flexibility and assessment at every step: outliers and edge cases may reveal interesting biological differences, rather than sample processing or technical errors. Many workflow systems can be used for either type, but we note cases where their properties facilitate one of these types over the other.

#### Using workflows without learning workflow syntax

While the benefits of executing an analysis within a data-centric workflow system are immense, the learning curve associated with command-line systems can be daunting. It is possible to obtain the benefits of workflow systems without learning new syntax. Websites like Galaxy, Cavatica, and EMBL-EBI MGnify offer online portals in which users build workflows around publicly-available or user-uploaded data [12,13,14]. On the command line, many research groups have used workflow systems to wrap one or many analysis steps (specified in an underlying workflow language) in a more user-friendly command-line application that accepts user input and executes the analysis. These pipeline applications allow users to take advantage of workflow software without needing to write the workflow syntax or manage software installation for each analysis step. Some examples include the nf-core RNA-seq pipeline [1,15], the PiGx genomic analysis toolkit [16], the ATLAS metagenome assembly and binning pipeline [17,18], the Sunbeam metagenome analysis pipeline [19,20], and two from our own lab, the dammit eukaryotic transcriptome annotation pipeline [21] and the elvers *de novo* transcriptome pipeline [22]. These pipeline applications typically execute a series of standard steps, but many provide varying degrees of customizability ranging from tool choice to parameter specification.

#### Choosing a workflow system

If your use case extends beyond these tools, there are several scriptable workflow systems that offer comparable benefits for carrying out your own data-intensive analyses. Each has it own strengths, meaning each workflow software will meet an individuals computing goals differently (see **Table 1**). Our lab has adopted Snakemake [23], in part due to its integration with Python, its flexibility for building and testing new analyses in different languages, and its intuitive integration with software management tools (described below). Snakemake and Nextflow [25] are commonly used for developing new research pipelines, where flexibility and iterative, branching development is a key feature. Common Workflow Language (CWL) and Workflow Description Language (WDL) are workflow specification formats that are more geared towards scalability, making them ideal for production-level pipelines with hundreds of thousands of samples [26]. WDL and CWL are commonly executed on platforms such as Terra [27] or Seven Bridges Platform [28]. Language-specific workflow systems, such as ROpenSci’s Drake [29], can take full advantage of the language’s internal data structures, and provide automation and reproducibility benefits for workflows executed primarily within the language ecosystem.

**Table 1:** Four of the most widely used bioinformatics workflow systems (2020), with links to documentation, example workflows, and general tutorials. In many cases, there may be tutorials online that are tailored for use cases in your field. All of these systems can interact with tools or tasks written in other languages and can function across cloud computing systems and high-performance computing clusters. Some can also import full workflows from other specification languages.

| Workflow System | Documentation | Example Workflow | Tutorial |
| --- | --- | --- | --- |
| Snakemake | https://snakemake.readthedocs.io/ | https://github.com/snakemake-workflows/chipseq | https://snakemake.readthedocs.io/en/stable/tutorial/tutorial.html |
| Nextflow | https://www.nextflow.io/ | https://github.com/nf-core/sarek | https://www.nextflow.io/docs/latest/getstarted.html |
| Common workflow language | https://www.commonwl.org/ | https://github.com/EBI-Metagenomics/pipeline-v5 | https://www.commonwl.org/user_guide/02-1st-example/index.html |
| Workflow description language | https://openwdl.org/ | https://github.com/gatk-workflows/gatk4-data-processing | https://support.terra.bio/hc/en-us/articles/360037127992-1-howto-Write-your-first-WDL-script-running-GATK-HaplotypeCaller |

The best workflow system to choose may be the one with a strong and accessible local or online community in your field, somewhat independent of your computational needs. The availability of field-specific data analysis code for reuse and modification can facilitate the adoption process, as can community support for new users. Fortunately, the standardized syntax required by workflow systems, combined with widespread adoption in the open science community, has resulted in a proliferation of open access workflow-system code for routine analysis steps [30,31]. At the same time, consensus approaches for data analysis are emerging, further encouraging reuse of existing code [32,33,34,35,36].

The Getting started developing workflows section contains strategies for modifying and developing workflows for your own analyses.

### Wrangling Scientific Software

Analysis workflows commonly rely on multiple software packages to generate final results. These tools are heterogeneous in nature: they are written by researchers working in different coding languages, with varied approaches to software design and optimization, and often for specific analysis goals. Each program has a number of other programs it depends upon to function (“dependencies”), and as software changes over time to meet research needs, the results may change, even when run with identical parameters. As a result, it is critical to take an organized approach to installing, managing, and keeping track of software and software versions. On many compute systems, system-wide software management is overseen by system administrators, who ensure commonly-used and requested software is installed into a “module” system available to all users. Unfortunately, this system limits software version transparency and does not lend itself well to exploring new workflows and software, as researchers do not have permission to install software themselves. To meet this need, most workflow managers integrate with software management systems that handle software installation, management, and packaging, alleviating problems that arise from complex dependencies and facilitating documentation of software versions. Software management systems range from lightweight systems that manage only the software and its dependencies, to heavyweight systems that control for all aspects of the runtime and operating system, ensuring 100% reproducibility of results across computational platforms and time.

On the lightweight end, the conda package manager has emerged as a leading software management solution for research workflows (**Figure 2**). Conda handles both cluster permission and version conflict issues with a user-based software environment system, and features a straightforward “recipe” system which simplifies the process of making new software installable (including simple management of versions and updates). These features have led to widespread adoption within the bioinformatics community: packages for new software become quickly available, and can be installed easily across platforms. However, conda does not completely isolate software installations and aims neither for bitwise reproducibility nor long-term archiving of install packages, meaning installations will not be completely reproducible over time. Heavyweight software management systems package not only the software of interest, but also the runtime environment information, with the goal of ensuring perfect reproducibility in software installation over time. Tools such as singularity and docker [3,11,37,38] wrap software environments in “containers” that capture and reproduce the runtime environment information. Container-based management is particularly useful for systems where some dependencies may not be installable by lightweight managers. However, software installation within these containers can be limited by similar reproducibility issues, including changes in dependency installations over time. “Functional package managers” such as GNU Guix and Nix strictly require all dependency and configuration details be encoded within each software package, providing the most comprehensively reproducible installations. These have begun to be integrated into some bioinformatic tools [16], but have a steeper learning curve for independent use. In addition, standard installation of these managers requires system-wide installation permissions, requiring assistance from system administrators on most high-performance computing systems.

**Figure 2:**
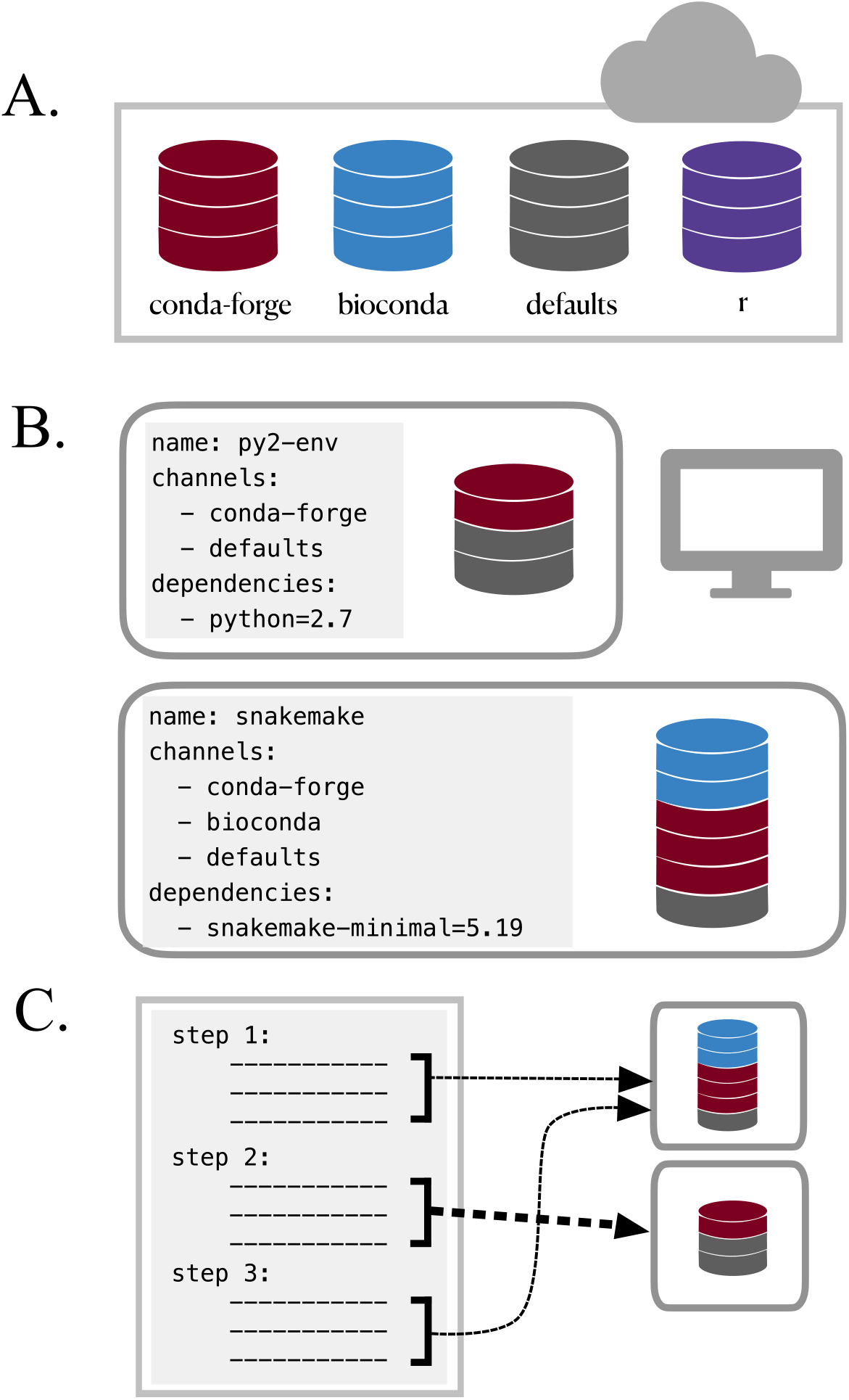
The conda package and environment manager simplifies software installation and management. **A. Conda Recipe Repositories:** Each program distributed via Conda has a “recipe” describing all software dependencies needed for installation using Conda (each of which must also be installable via Conda). Recipes are stored and managed in the cloud in separate “channels”, some of which specialize in particular fields or languages (e.g. the “bioconda” channel specializes in bioinformatic software, while the “conda-forge” channel is a more general effort to provide and maintain standardized conda packages for a wide range of software) [11]. **B. Use Conda Environments to Avoid Installation Conflicts:** Conda does not require root privileges for software installation, thus enabling use by researchers working on shared cluster systems. However, even user-based software installation can encounter dependency conflicts. For example, you might need to use python2 to install and run a program (e.g. older scripts written by members of your lab), while also using snakemake to execute your workflows (requires python>=3.5). By installing each program into an isolated “environment” that contains only the software required to run that program, you can ensure all programs used throughout your analysis will run without issue. Using small, separate environments for your software, specifying the desired software version, and building many simple environments to accommodate different steps in your workflow is critical for reducing the amount of time it takes conda to resolve dependency conflicts between different software tools (“solve” an environment). Conda virtual environments can be created and installed either on the command line, or via an environment YAML file, as shown. In this case, the environment file also specifies which conda channels to search and download programs from. When specified in a YAML file, conda environments are easily transferable between computers and operating systems. Broad community adoption has resulted in a proliferation of both conda-installable scientific software and tools that leverage conda installation specifications. For example, the Mamba package manager is an open source reimplementation of the conda manager that can install conda-style environments with increased efficiency [39]. The BioContainers Registry is a project that automatically builds and distributes docker and singularity containers for bioinformatics software packages using each package’s conda installation recipe [40].

### Getting started with software management

#### Using software without learning software management systems

First, there are a number of ways to test software before needing to worry about installation. Some software packages are available as web-based tools and through a series of data upload and parameter specifications, allow the user to interact with a tool that is running on a back-end server. Integrated development environments (IDE) like PyCharm and RStudio can manage software installation for language-specific tools, and can be very helpful when writing analysis code. While these approaches do not integrate into reproducible workflows, they may be ideal for testing a tool to determine whether it is useful for your data before integration in your analysis.

#### Choosing a software management system

It is important to balance the time needed to learn to properly use a software management system with the needs of both the project and the researchers. Software management systems with large learning curves are less likely to be widely adopted among researchers with a mix of biological and computational backgrounds. In our experience, software management with conda nicely balances reproducibility with flexibility and ease of use. These trade-offs are best for research workflows under active development, where flexible software installation solutions that enable new analysis explorations or regular tool updates are critical. For production workflows that require maximal reproducibility, it is worth the larger investment required to use heavyweight systems. This is particularly true for advanced users who can more easily navigate the steps required for utilizing these tools. Container-based software installation via docker and singularity are common for production-level workflows, and Guix and Nix-based solutions are gaining traction. Importantly, the needs and constraints of a project can evolve over time, as may the system of choice.

#### Integrating software management within workflows

Workflow systems provide seamless integration with a number of software management tools. Each workflow system requires different specification for initiation of software management, but typically requires about one additional line of code per step that requires the use of software. If the software management tool is installed locally, the workflow will automatically download and install the specified environment or container and use it for specified step.

In our experience, the complete solution for using scientific software involves a combination of approaches. Interactive and exploratory analyses conducted in IDEs and jupyter notebooks (usually with local software installation with conda) are useful for developing an analysis strategy and creating an initial workflow. This is then followed by workflow-integrated software management via conda, singularity, or nixOS for executing the resulting workflow on many samples. This process not linear: we often cycle between exploratory testing and automation as we iteratively extend our analyses.

## Workflow-Based Project Management

Project management, the strategies and decisions used to keep a project organized, documented, functional, and shareable, is foundational to any research program. Clear organization and management is a learned skill that takes time to implement. Workflow systems simplify and improve computational project management, but even workflows that are fully specified in workflow systems require additional investment to stay organized, documented, and backed up.

### Systematically document your workflows

Pervasive documentation provides indispensable context for biological insights derived from an analysis, facilitates transparency in research, and increases reusability of the analysis code. Good documentation covers all aspects of a project, including file and results organization, clear and commented code, and accompanying explanatory documents for design decisions and metadata. Workflow systems facilitate building this documentation, as each analysis step (with chosen parameters) and the links between those steps are completely specified within the workflow syntax. This feature streamlines code documentation, particularly if you include as much of the analysis as possible within the automated workflow framework. Outside of the analysis itself, applying consistent organizational design can capitalize on the structure and automation provided by workflows to simplify the generation of quality documentation for all aspects of your project. Below, we discuss project management strategies for building reproducible workflow-enabled biological analyses.

### Use consistent, self-documenting names

Using consistent and descriptive identifiers for your files, scripts, variables, workflows, projects, and even manuscripts helps keep your projects organized and interpretable for yourself and collaborators. For workflow systems, this strategy can be implemented by tagging output files with a descriptive identifier for each analysis step, either in the filename or by placing output files within a descriptive output folder. For example, the file shown in **Figure 3** has been preprocessed with a quality control trimming step. For large workflows, placing results from each step of your analysis in isolated, descriptive folders can be essential for keeping your project workspace clean and organized.

**Figure 3:**
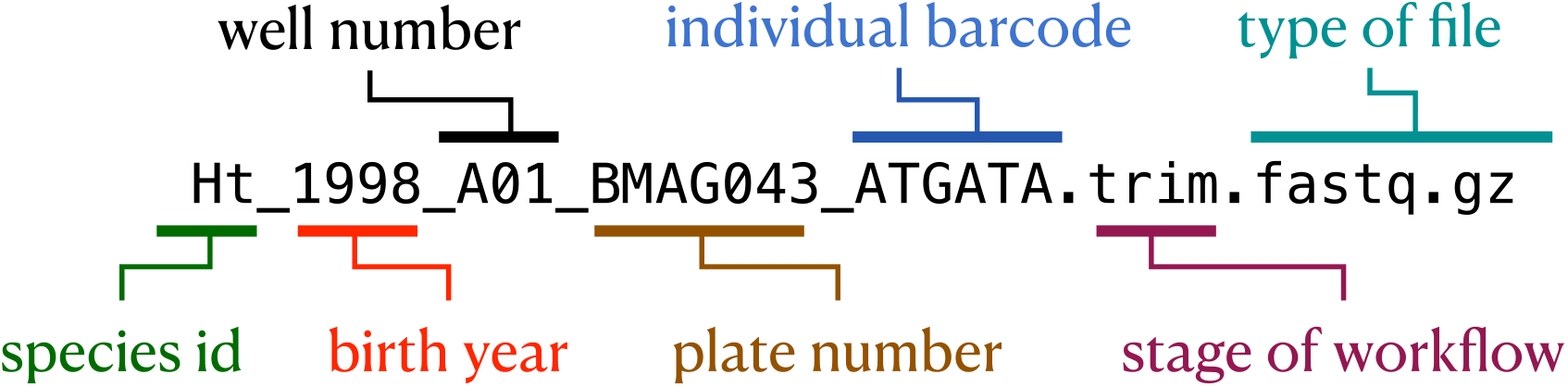
Consistent and informative file naming improves organization and interpretability. For ease of grouping and referring to input files, it is useful to keep unique sample identification in the filename, often with a metadata file explaining the meaning of each unique descriptor. For analysis scripts, it can help to implement a numbering scheme, where the name of first file in the analysis begins with “00”, the next with “01”, etc. For output files, it can help to add a short, unique identifier to output files processed with each analysis step. This particular file is a RAD sequencing fastq file of a fish species that has been preprocessed with a fastq quality trimming tool.

### Store workflow metadata with the workflow

Developing biological analysis workflows can involve hundreds of small decisions: What parameters work best for each step? Why did you use a certain reference file for annotation as compared with other available files? How did you finally manage to get around the program or installation error? All of these pieces of information contextualize your results and may be helpful when sharing your findings. Keeping information about these decisions in an intuitive and easily accessible place helps you find it when you need it. To capitalize on the utility of version control systems described below, it is most useful to store this information in plain text files. Each main directory of a project should include notes on the data or scripts contained within, so that a collaborator could look into the directory and understand what to find there (especially since that “collaborator” is likely to be you, a few months from now!). Code itself can contain documentation - you can include comments with the reasoning behind algorithm choice or include a link to online documentation or solution that helped you decide how to shape your differential expression analysis. Larger pieces of information can be kept in “README” or notes documents kept alongside your code and other documents. For example, a GitHub repository documenting the reanalysis of the Marine Microbial Eukaryote Transcriptome Sequencing Project uses a README alongside the code to document the workflow and digital object identifiers for data products [41,42]. While this particular strategy cannot be automated, it is critical for interpreting the final results of your workflow.

### Document data and analysis exploration using computational notebooks

Computational notebooks allow users to combine narrative, code, and code output (e.g. visualizations) in a single location, enabling the user to conduct analysis and visually assess the results in a single file (see **Figure 4**). These notebooks allow for fully documented iterative analysis development, and are particularly useful for data exploration and developing visualizations prior to integration into a workflow or as a report generated by a workflow that can be shared with collaborators.

**Figure 4:**
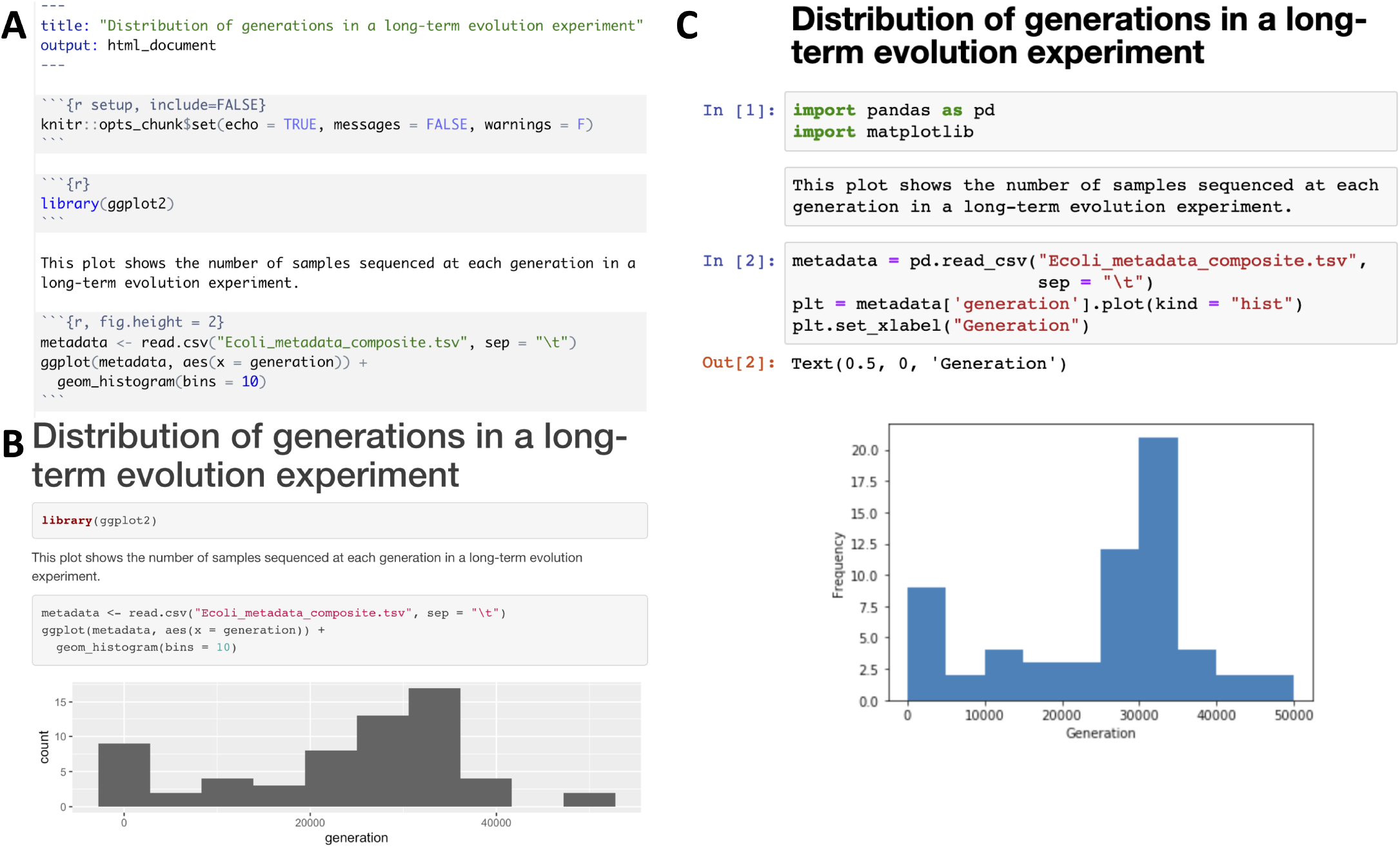
Examples of computational notebooks. Computational notebooks allow the user to mix text, code, and results in one document. **Panel A.** shows an RMarkdown document viewed in the RStudio integrated development environment, while **Panel B.** shows a rendered HTML file produced by knitting the RMarkdown document [43]. **Panel C.** shows a Jupyter Notebook, where code, text, and results are rendered inline as each code chunk is executed [44]. The second grey chunk is a raw Markdown chunk with text that will be rendered inline when executed. Both notebooks generate a histogram of a metadata feature, number of generations, from a long-term evolution experiment with *Escherichia coli* [45]. Computational notebooks facilitate sharing by packaging narrative, code, and visualizations together. Sharing can be enhanced further by packaging computational notebooks with tools like Binder [46]. Binder builds an executable environment (capable of running RStudio and Jupyter notebooks) out of a GitHub repository using package management systems and docker to build reproducible and executable software environments as specified in the repository. Binders can be shared with collaborators (or students in a classroom setting), and analysis and visualization can be ephemerally reproduced or altered from the code provided in computational notebooks.

### Visualize your workflow

Visual representations can help illustrate the connections in a workflow and improve the readability and reproducibility of your project. At the highest level, flowcharts that detail relationships between steps of a workflow can help provide big-picture clarification, especially when the pipeline is complicated. For individual steps, a graphical representation of the output can show the status of the project or provide insight on additional analyses that should be added. For example, **Figure 5** exhibits a modified Snakemake workflow visualization from an RNA-seq quantification pipeline [47].

**Figure 5:**
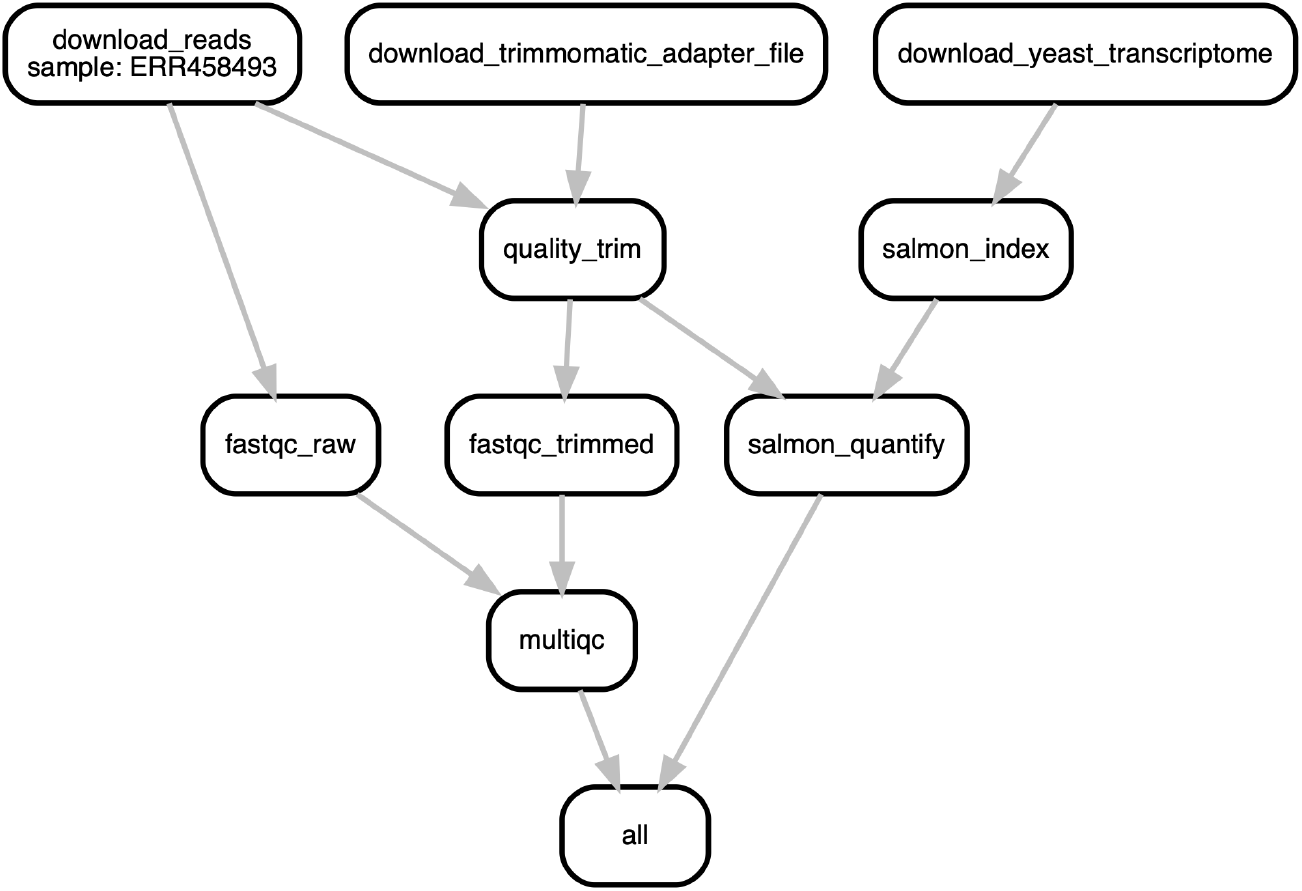
A directed acyclic graph (DAG) that illustrates connections between all steps of a sequencing data analysis workflow. Each box represents a step in the workflow, while lines connect sequential steps. The DAG shown in this figure illustrates a real bioinformatics workflow for RNA-seq quantification was generated by modifying the default Snakemake workflow DAG. This example of an initial workflow used only to quality control and then quantify one FASTQ file against a transcriptome more than doubles the amount of files in a project. When the number of steps are expanded to carry out a full research analysis and the number of initial input files are increased, a workflow can generate hundreds to thousands of intermediate files. Fortunately, workflow system coordination alleviates the need for a user to directly manage file interdependencies. For a larger analysis DAG, see [48]

### Version control your project

As your project develops, version control allows you to keep track of changes over time. You may already do this in some ways, perhaps with frequent hard drive backups or by manually saving different versions of the same file - e.g. by appending the date to a script name or appending “version_1” or “version_FINAL” to a manuscript draft. For computational workflows, using version control systems such as Git or Mercurial can be used to keep track of all changes over time, even across multiple systems, scripting languages, and project contributors (see **Figure 6**). If a key piece of a workflow inexplicably stops working, consistent version control can allow you to rewind in time and identify differences from when the pipeline worked to when it stopped working. Backing up your version controlled analysis in an online repository such as GitHub, GitLab, or Bitbucket provides critical insurance as you iteratively modify and develop your workflow.

**Figure 6:**
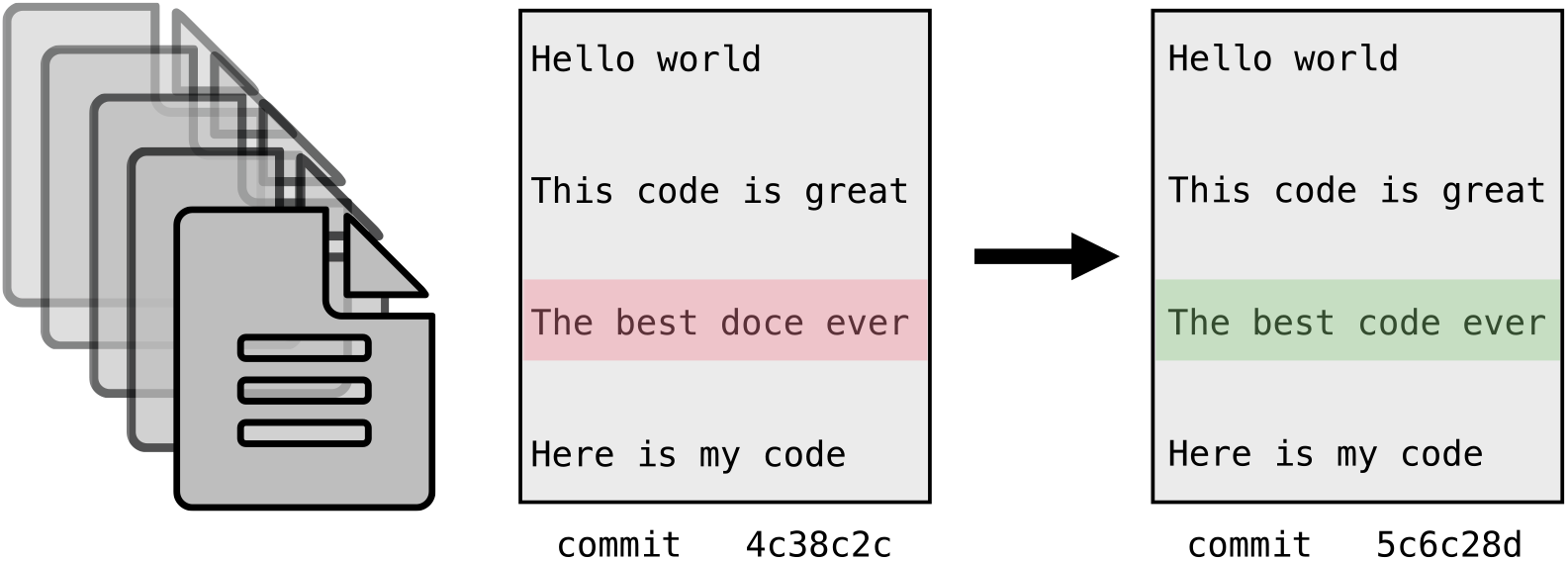
Version Control. Version control systems (e.g. Git, Mercurial) work by storing incremental differences in files from one saved version (“commit”) to the next. To visualize the differences between each version, text editors such as Atom and online services such as GitHub, GitLab and Bitbucket use red highlighting to denote deletions, and green highlighting to denote additions. In this trivial example, a typo in version 1 (in red) was corrected in version 2 (in green). These systems are extremely useful for code and manuscript development, as it is possible to return to the snapshot of any saved version. This means that version control systems save you from accidental deletions, preserve code you thought you no longer needed and preserve a record of project changes over time.

When combined with online backups, version control systems also facilitate code and data availability and reproducibility for publication. For example, to preserve the version of code that produced published results, you can create a “release”: a snapshot of the current code and files in a GitHub repository. You can then generate a digital object identifier (DOI) for that release using a permanent documentation service such as Zenodo ([49]) and make it available to reviewers and beyond (see “sharing” section, below).

### Share your workflow and analysis code

Sharing your workflow code with collaborators, peer reviewers, and scientists seeking to use a similar method can foster discussion and review of your analysis. Sticking to a clear documentation strategy, using a version control system, and packaging your code in notebooks or as a workflow prepare them to be easily shared with others. To go one step further, you can package your code with tools like Binder, ReproZip, or Whole Tale, or make interactive visualizations with tools like Shiny apps or Plotly. These approaches let others run the code on cloud computers in environments identical to those in which the original computation was performed (**Figure 4**, **Figure 7**) [46,50,51]. These tools substantially reduce overhead associated with interacting with code and data, and in doing so, make it fast and easy to rerun portions of the analysis, check accuracy, or even tweak the analysis to produce new results. If you also share your code and workflows publicly, you will also help contribute to the growing resources for open workflow-enabled biological research.

**Figure 7:**
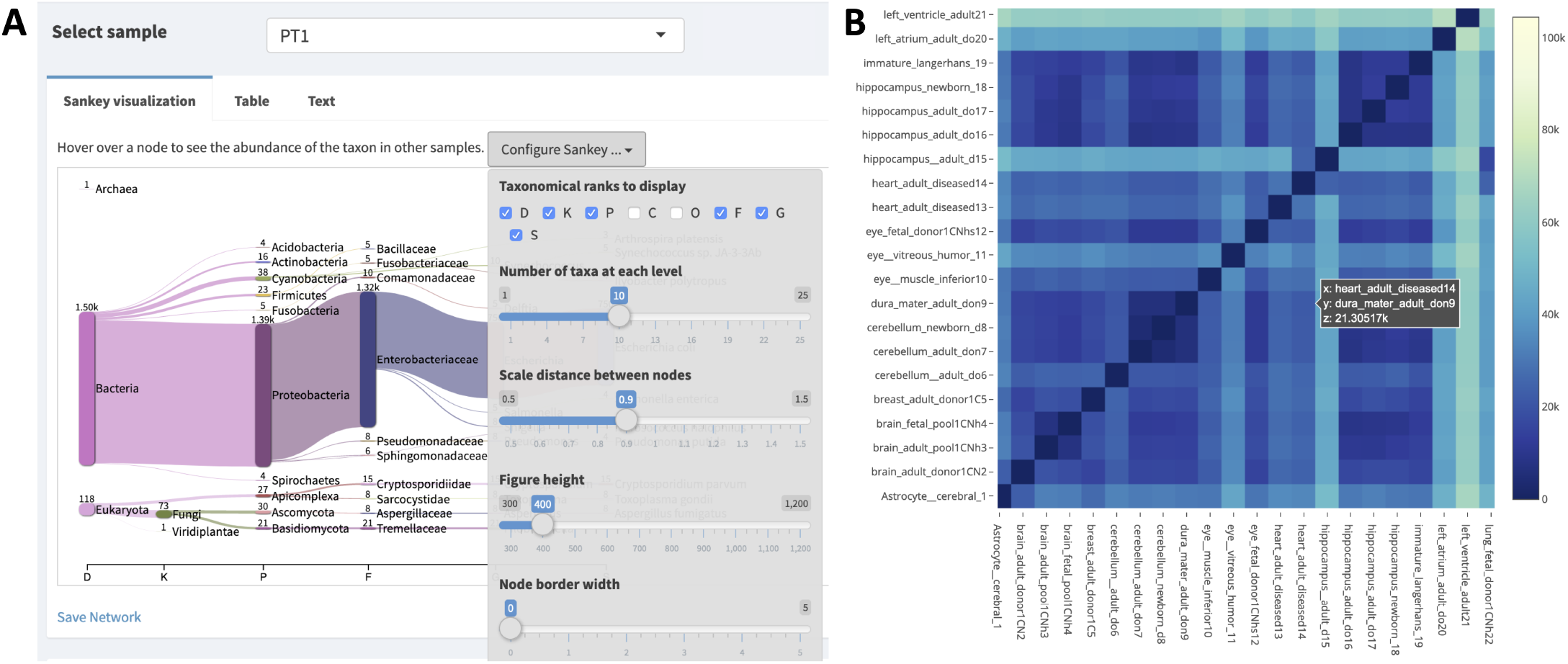
Interactive visualizations facilitate sharing and repeatability. **A.** Interactive visualization dashboard in the Pavian Shiny app for metagenomic analysis [52,53]. Shiny allows you to build interactive web pages using R code. Data is manipulated by R code in real-time in a web page, producing analysis and visualizations of a data set. Shiny apps can contain user-specifiable parameters, allowing a user to control visualizations or analyses. As seen above, sample “PT1” is selected, and taxonomic ranks class and order are excluded. Shiny apps allow collaborators who may or may not know R to modify R visualizations to fit their interests. **B.** Plotly heatmap of transcriptional profiling in human brain samples [54]. Hovering over a cell in the heatmap displays the sample names from the x and y axis, as well as the intensity value. Plotting tools like plotly and vega-lite produce single interactive plots that can be shared with collaborators or integrated into websites [55,56]. Interactive visualizations are also helpful in exploratory data analysis.

### Getting started developing workflows

In our experience, the best way to have your workflow system work *for* you is to include as much of your analysis as possible within the automated workflow framework, use self-documenting names, include analysis visualizations, and keep rigorous documentation alongside your workflow that enables you to understand each decision and entirely reproduce any manual steps. Some of the tools discussed above will inevitably change over time, but these principles apply broadly and will help you design clear, well-documented, and reproducible analyses. Ultimately, you will need to experiment with strategies that work for you – what is most important is to develop a clear set of strategies and implement them tenaciously. Below, we provide a few practical strategies to try as you begin developing your own workflows.

#### Start with working code

When building a workflow for the first time, start from working examples provided as part of the tool documentation or otherwise available online. This functioning example code then provides a reliable workflow framework free of syntax errors which you can customize for your data without the overhead of generating correct workflow syntax from scratch. Be sure to run this analysis on provided test data, if available, to ensure the tools, and command line syntax function at a basic level. **Table 1** provides links to official repositories containing tutorials and example biological analysis workflows, and workflow tutorials and code sharing websites like GitHub, GitLab, and Bitbucket have many publicly available workflows for other analyses. If a workflow is available through Binder, you can test and experiment with workflow modification on Binder’s cloud system without needing to install a workflow manager or software management tool on your local compute system [46].

#### Test with subsampled data

Once you have working workflow syntax, test the step on your own data or public data related to your species or condition of interest. First, create a subsampled dataset that you can use to test your entire analysis workflow. This set will save time, energy, and computational resources throughout workflow development. If working with FASTQ data, a straightforward way to generate a small test set is to subsample the first million lines of a file (first 250k reads):

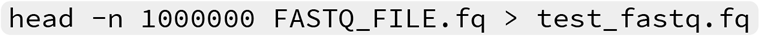

While there are many more sophisticated ways to subsample reads, this technique should be sufficient for testing each step of a most workflows prior to running your full dataset. In specific cases, such as eukaryotic genome assembly, you may need to be more intentional with how you subsample reads and how much sample data you use as a test set.

#### Document your process

Document your changes, explorations, and errors as you develop. We recommend using the Markdown language so your documentation is in plain text (to facilitate version control), but can still include helpful visual headings, code formatting, and embedded images. Markdown editors with visual previewing, such as HackMD, can greatly facilitate notetaking, and Markdown documents are visually rendered properly within your online version control backups on services such as GitHub [57].

#### Develop your workflow

From your working code, iteratively modify and add workflow steps to meet your data analysis needs. This strategy allows you to find and fix mistakes on small sections of the workflow. Periodically clean your output directory and rerun the entire workflow, to ensure all steps are fully interoperable (using small test data will improve the efficiency of this step!). If possible, using mock or control datasets can help you verify that the analysis you are building actually returns correct biological results. Tutorials and tool documentation are useful companions during development; as with any language, remembering workflow-specific syntax takes time and practice.

#### Assess your results

Evaluate your workflow results as you go. Consider what aspects (e.g. tool choice, program parameters) can be evaluated rigorously, and assess each step for expected behavior. Other aspects (e.g. filtering metadata, joining results across programs or analysis, software and workflow bugs) will be more difficult to evaluate. Wherever possible, set up positive and negative controls to ensure your analysis is performing the desired analysis properly. Once you’re certain an analysis is executing as designed, tracking down unusual results may reveal interesting biological differences.

#### Back up early and often

As you write new code, back up your changes in an online repository such as GitHub, GitLab, or Bitbucket. These services support both drag-and-drop and command line interaction.

#### Scale up your workflow

Bioinformatic tools vary in the resources they require: some analysis steps are compute-intensive, other steps are memory intensive, and still others will have large intermediate storage needs. If using high-performance computing system or the cloud, you will need to request resources for running your pipeline, often provided as a simultaneous execution limit or purchased by your research group on a cost-per-compute basis. Workflow systems provide built-in tools to monitor resource usage for each step. Running a complete workflow on a single sample with resource monitoring enabled generates an estimate of computational resources needed for each step. These estimates can be used to set appropriate resource limits for each step when executing the workflow on your remaining samples.

#### Find a community and ask for help when you need it

Local and online users groups are helpful communities when learning a workflow language. When you are first learning, help from more advanced users can save you hours of frustration. After you’ve progressed, providing that same help to new users can help you cement the syntax in your mind and tackle more advanced uses. Data-centric workflow systems have been enthusiastically adopted by the open science community, and as a consequence, there is a critical mass of tutorials and open access code, as well as code discussion on forums and via social media, particularly Twitter. Post in the relevant workflow forums when you have hit a stopping point you are unable to work through. Be respectful of people’s time and energy and be sure to include appropriate details important to your problem (see Strategic troubleshooting section).

## Data and resource management for workflow-enabled biology

Advancements in sequencing technologies have greatly increased the volume of data available for biological query [58]. Workflow systems, by virtue of automating many of the time-intensive project management steps traditionally required for data-intensive biology, can increase our capacity for data analysis. However, conducting biological analyses at this scale requires a coordinated approach to data and computational resource management. Below, we provide recommendations for data acquisition, management, and quality control that have become especially important as the volume of data has increased. Finally, we discuss securing and managing appropriate computational resources for the scale of your project.

### Managing large-scale datasets

Experimental design, finding or generating data, and quality control are quintessential parts of data intensive biology. There is no substitute for taking the time to properly design your analysis, identify appropriate data, and conduct sanity checks on your files. While these tasks are not automatable, many tools and databases can aid in these processes.

### Look for appropriate publicly-available data

With vast amounts of sequencing data already available in public repositories, it is often possible to begin investigating your research question by seeking out publicly available data. In some cases, these data will be sufficient to conduct your entire analysis. In others cases, particularly for biologists conducting novel experiments, these data can inform decisions about sequencing type, depth, and replication, and can help uncover potential pitfalls before they cost valuable time and resources.

Most journals now require data for all manuscripts to be made accessible, either at publication or after a short moratorium. Further, the FAIR (findable, accessible, interoperable, reusable) data movement has improved the data sharing ecosystem for data-intensive biology [59,60,61,62,63,64,64,65]. You can find relevant sequencing data either by starting from the “data accessibility” sections of papers relevant to your research or by directly searching for your organism, environment, or treatment of choice in public data portals and repositories. The International Nucleotide Sequence Database Collaboration (INSDC), which includes the Sequence Read Archive (SRA), European Nucleotide Archive (ENA), and DataBank of Japan (DDBJ) is the largest repository for raw sequencing data, but no longer accepts sequencing data from large consortia projects [66]. These data are instead hosted in consortia-specific databases, which may require some domain-specific knowledge for identifying relevant datasets and have unique download and authentication protocols. For example, raw data from the Tara Oceans expedition is hosted by the Tara Ocean Foundation [67]. Additional curated databases focus on processed data instead, such as gene expression in the Gene Expression Omnibus (GEO) [68]. Organism-specific databases such as **Wormbase** (*Caenorhabditis elegans*) specialize on curating and integrating sequencing and other data associated with a model organism [69]. Finally, rather than focusing on certain data types or organisms, some repositories are designed to hold any data and metadata associated with a specific project or manuscript (e.g. Open Science Framework, Dryad, Zenodo [70]).

### Consider analysis when generating your own data

If generating your own data, proper experimental design and planning are essential. For cost-intensive sequencing data, there are a range of decisions about experimental design and sequencing (including sequencing type, sequencing depth per sample, and biological replication) that impact your ability to properly address your research question. Conducting discussions with experienced bioinformaticians and statisticians, *prior to beginning your experiments* if possible, is the best way to ensure you will have sufficient statistical power to detect effects. These considerations will be different for different types of sequence analysis. To aid in early project planning, we have curated a series of domain-specific references that may be useful as you go about designing your experiment (see **Table 2**). Given the resources invested in collecting samples for sequencing, it’s important to build in a buffer to preserve your experimental design in the face of unexpected laboratory or technical issues. Once generated, it is always a good idea to have multiple independent backups of raw sequencing data, as it typically cannot be easily regenerated if lost to computer failure or other unforeseeable events.

**Table 2:** References for experimental design and considerations for common sequencing chemistries.

| Sequencing type | Resources |
| --- | --- |
| RNA-sequencing | [32,71,72] |
| Metagenomic sequencing | [33,73,74] |
| Amplicon sequencing | [75,76,77] |
| Microbial isolate sequencing | [78] |
| Eukaryotic genome sequencing | [79,80,81,82] |
| Whole-genome resequencing | [83] |
| RAD-sequencing | [84,84,85,86,87,88] |
| single cell RNA-seq | [89,90] |

As your experiment progresses, keep track of as much information as possible: dates and times of sample collection, storage, and extraction, sample names, aberrations that occurred during collection, kit lot used for extraction, and any other sample and sequencing measurements you might be able to obtain (temperature, location, metabolite concentration, name of collector, well number, plate number, machine your data was sequenced, on etc). This metadata allows you to keep track of your samples, to control for batch effects that may arise from unintended batching during sampling or experimental procedures and makes the data you collect reusable for future applications and analysis by yourself and others. Wherever possible, follow the standard guidelines for formatting metadata for scientific computing to limit downstream processing and simplify analyses requiring these metadata (see: [10]). We have focused here on sequencing data; for data management over long-term ecological studies, we recommend [91].

### Getting started with sequencing data

#### Protect valuable data

Aside from the code itself, raw data are the most important files associated with a workflow, as they cannot be regenerated if accidentally altered or deleted. Keeping a read-only copy of raw data alongside a workflow as well multiple backups protects your data from accidents and computer failure. This also removes the imperative of storing intermediate files as these can be easily regenerated by the workflow.

When sharing or storing files and results, data version control can keep track of differences in files such as changes from tool parameters or versions. The version control tools discussed in the Workflow-based project management section are primarily designed to handle small files, but GitHub provides support for Git Large File Storage (LFS), and repositories such as the Open Science Framework (OSF), Figshare, Zenodo, and Dryad can be used for storing larger files and datasets [49,70,92,93,94].

In addition to providing version control for projects and datasets, these tools also facilitate sharing and attribution by enabling generation of digital object identifiers (doi) for datasets, figures, presentations, code, and preprints. As free tools often limit the size of files that can be stored, a number of cloud backup and storage services are also available for purchase or via university contract, including Google Drive, Box, Dropbox, Amazon Web Services, and Backblaze. Full computer backups can be conducted to these storage locations with tools like rclone [95].

#### Ensure data integrity during transfers

If you’re working with publicly-available data, you may be able to work on a compute system where the data are already available, circumventing time and effort required for downloading and moving the data. Databases such as the Sequence Read Archive (SRA) are now available on commercial cloud computing systems, and open source projects such as Galaxy enable working with SRA sequence files directly from a web browser [12,96]. Ongoing projects such as the NIH Common Fund Data Ecosystem aim to develop a data portal to make NIH Common Fund data, including biomedical sequencing data, more findable, accessible, interoperable, and reusable (FAIR).

In most cases, you’ll still need to transfer some data - either downloading raw data or transferring important intermediate and results files for backup and sharing (or both). Transferring compressed files (gzip, bzip2, BAM/CRAM, etc.) can improve transfer speed and save space, and checksums can be used to to ensure file integrity after transfer (see **Figure 8**).

**Figure 8:**
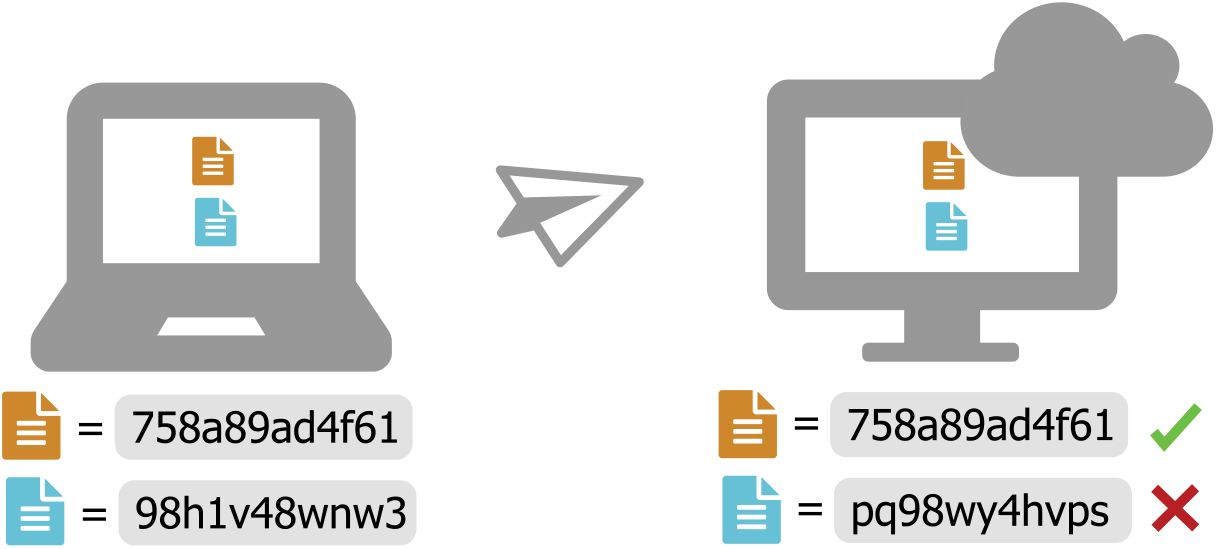
Use Checksums to ensure file integrity. Checksum programs (e.g. md5, sha256) encode file size and content in a single value known as a “checksum”. For any given file, this value will be identical across platforms when calculated using the same checksum program. When transferring files, calculate the value of the checksum prior to transfer, and then again after transfer. If the value is not identical, there was an error introduced during transfer (e.g. file truncation, etc). Checksums are often provided alongside publicly available files, so that you can verify proper download. Tools like rsync and rclone that automate file transfers use checksums internally to verify that files were transferred properly, and some GUI file transfer tools (e.g. Cyberduck) can assess checksums when they are provided [95]. If you generated your own data and receieved sequencing files from a sequencing center, be certain you also receive a checksum for each of your files to ensure they download properly.

#### Perform quality control at every step

The quality of your input data has a major impact on the quality of the output results, no matter whether your workflow analyzes six samples or six hundred. Assessing data at every analysis step can reveal problems and errors early, before they waste valuable time and resources. Using quality control tools that provide metrics and visualizations can help you assess your datasets, particularly as the size of your input data scales up. However, data from different species or sequencing types can produce anomalous quality control results. You are ultimately the single most effective quality control tool that you have, so it is important to critically assess each metric to determine those that are relevant for your particular data.

##### Look at your files

Quality control can be as simple as looking at the first few and last few lines of input and output data files, or checking the size of those files (see **Table 3**). To develop an intuition for what proper inputs and outputs look like for a given tool, it is often helpful to first run the test example or data that is packaged with the software. Comparing these input and output file formats to your own data can help identify and address inconsistencies.

**Table 3:** Some commands to quickly explore the contents of a file. These commands can be used on Unix and Linux operating systems to detect common formatting problems or other abnormalities.

| command | function | example |
| --- | --- | --- |
| ls -lh | list files with information in a human-readable format | ls -lh *fastq.gz |
| head | print the first 6 lines of a file to standard out | head samples.csv |
| tail | print the last 6 lines of a file to standard out | tail samples.csv |
| less | show the contents of a file in a scrollable screen | less samples.csv |
| zless | show the contents of a gzipped file in a scrollable screen | zless sample1.fastq.gz |
| wc -l | count the number of lines in a file | wc -l ecoli.fasta |
| cat | print a file to standard out | cat samples.csv |
| grep | find matching text and print the line to standard out | grep ">" ecoli.fasta |
| cut | cut columns from a table | cut -d"," -f1 samples.csv |

##### Visualize your data

Visualization is another powerful way to pick out unusual or unexpected patterns. Although large abnormalities may be clear from looking at files, others may be small and difficult to find. Visualizing raw sequencing data with FastQC (**Figure 9A**) and processed sequencing data with tools like the Integrative Genome Viewer and plotting tabular results files using python or R can make aberrant or inconsistent results easier to track down [98,99].

**Figure 9:**
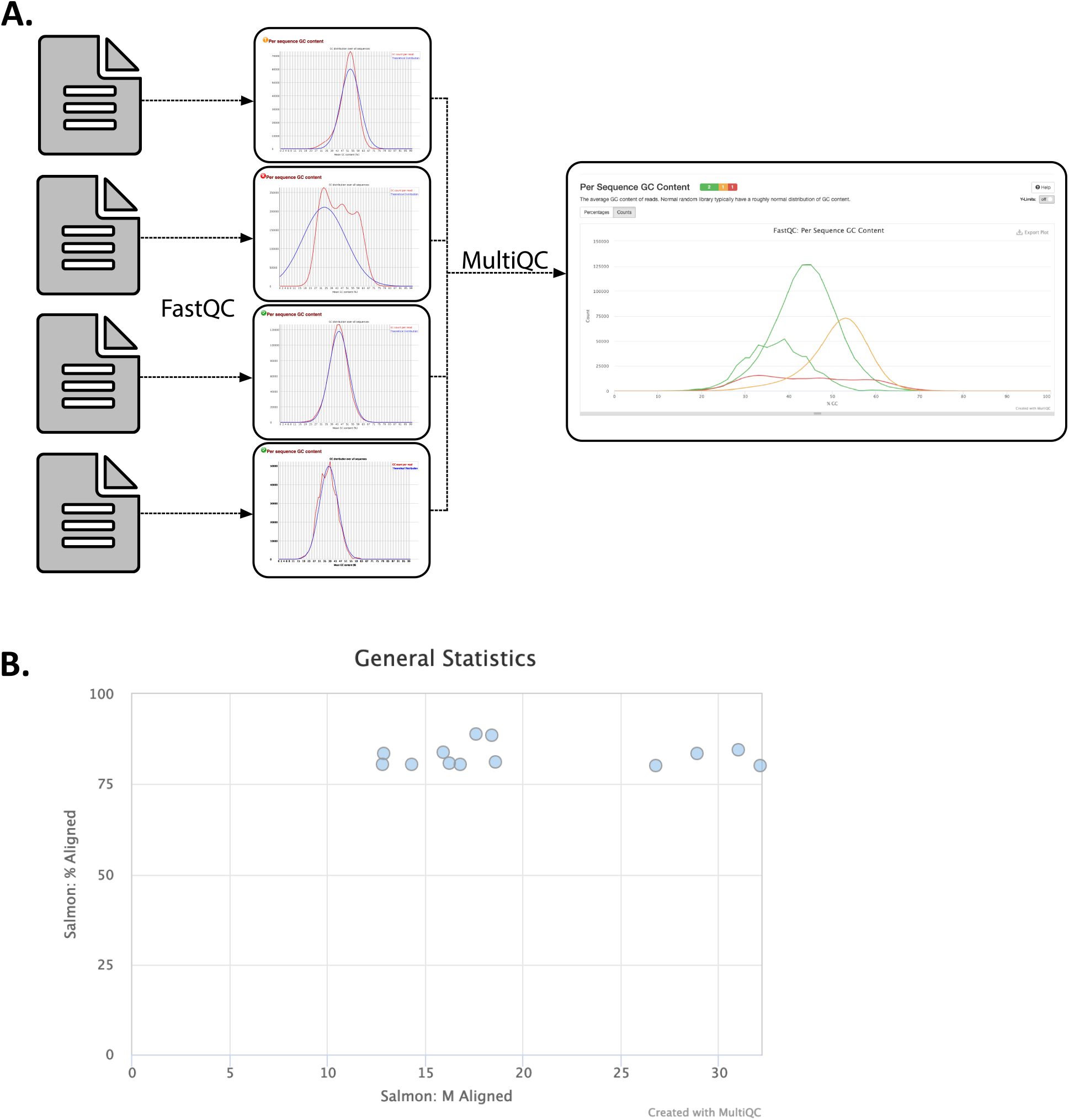
Visualizations produced by MultiQC. MultiQC finds and automatically parses log files from other tools and generates a combined report and parsed data tables that include all samples. MultiQC currently supports 88 tools. **A.** MultiQC summary of FastQC Per Sequence GC Content for 1905 metagenome samples. FastQC provides quality control measurements and visualizations for raw sequencing data from a single sample, and is a near-universal first step in sequencing data analysis because of the insights it provides [98,99]. FastQC measures and summarizes 10 quality metrics and provides recommendations for whether an individual sample is within an acceptable quality range. Not all metrics readily apply to all sequencing data types. For example, while multiple GC peaks might be concerning in whole genome sequencing of a bacterial isolate, we would expect a non-normal distribution for some metagenome samples that contain organisms with diverse GC content. Samples like this can be seen in red in this figure. **B.** MultiQC summary of Salmon *quant* reads mapped per sample for RNA-seq samples [100]. In this figure, we see that MultiQC summarizes the number of reads mapped and percent of reads mapped, two values that are reported in the Salmon log files.

##### Pay attention to warnings and log files

Many tools generate log files or messages while running. These files contain information about the quantity, quality, and results from the run, or error messages about why a run failed. Inspecting these files can be helpful to make sure tools ran properly and consistently, or to debug failed runs. Parsing and visualizing log files with a tool like MultiQC can improve interpretability of program-specific log files (**Figure 9**[101]).

##### Look for common biases in sequencing data

Biases in sequencing data originate from experimental design, methodology, sequencing chemistry, or workflows, and are helpful to target specifically with quality control measures. The exact biases in a specific data set or workflow will vary greatly between experiments so it is important to understand the sequencing method you have chosen and incorporate appropriate filtration steps into your workflow. For example, PCR duplicates can cause problems in libraries that underwent an amplification step, and often need to be removed prior to downstream analysis [102,103,104,105,106].

##### Check for contamination

Contamination can arise during sample collection, nucleotide extraction, library preparation, or through sequencing spike-ins like PhiX, and could change data interpretation if not removed [107,108,109]. Libraries sequenced with high concentrations of free adapters or with low concentration samples may have increased barcode hopping, leading to contamination between samples [110].

##### Consider the costs and benefits of stringent quality control for your data

Good quality data is essential for good downstream analysis. However, stringent quality control can sometimes do more harm than good. For example, depending on sequencing depth, stringent quality trimming of RNA-sequencing data may reduce isoform discovery [111]. To determine what issues are most likely to plague your specific data set, it can be helpful to find recent publications using a similar experimental design, or to speak with experts at a sequencing core.

Because sequencing data and applications are so diverse, there is no one-size-fits-all solution for quality control. It is important to think critically about the patterns you expect to see given your data and your biological problem, and consult with technical experts whenever possible.

### Securing and managing appropriate computational resources

Sequence analysis requires access to computing systems with adequate storage and analysis power for your data. For some smaller-scale datasets, local desktop or even laptop systems can be sufficient, especially if using tools that implement data-reduction strategies such as minhashing [112]. However, larger projects require additional computing power, or may be restricted to certain operating systems (e.g. linux). For these projects, solutions range from research-focused high performance computing systems to research-integrated commercial analysis platforms. Both research-only and and commercial clusters provide avenues for research and educational proposals to enable access to their computing resources (see **Table 4**). In preparing for data analysis, be sure to allocate sufficient computational resources and funding for storage and analysis, including large intermediate files and resources required for personnel training. Note that workflow systems can greatly facilitate faithful execution of your analysis across the range of computational resources available to you, including distribution across cloud computing systems.

**Table 4:** Computing Resources. Bioinformatic projects often require additional computing resources. If a local or university-run high-performance computing cluster is not available, computing resources are available via a number of grant-based or commercial providers.

| Provider | Access Model | Restrictions |
| --- | --- | --- |
| Amazon Web Services | Paid |  |
| Bionimbus Protected Data Cloud | Research allocation | users with eRA commons account |
| Cyverse Atmosphere | Free with limits | storage and compute hours |
| EGI federated cloud | Access by contact | European partner countries |
| Galaxy | Free with storage limits | data storage limits |
| Google Cloud Platform | Paid |  |
| Google Colab | Free | computational notebooks, no resource guarantees |
| Microsoft Azure | Paid |  |
| NSF XSEDE | Research allocation | USA researchers or collaborators |
| Open Science Data Cloud | Research allocation |  |
| Wasabi | Paid | data storage solution only |

### Getting started with resource management

As the scale of data increases, the resources required for analysis can balloon. Bioinformatic workflows can be long-running, require high-memory systems, or involve intensive file manipulation. Some of the strategies below may help you manage computational resources for your project.

#### Apply for research units if eligible

There are a number of cloud computing services that offer grants providing computing resources to data-intensive researchers (**Table 4**). In some cases, the resources provided may be sufficient to cover your entire analysis.

#### Develop on a local computer when possible

Since workflows transfer easily across systems, it can be useful to develop individual analysis steps on a local laptop. If the analysis tool will run on your local system, test the step with subsampled data, such as that created in the Getting started developing workflows section. Once working, the new workflow component can be run at scale on a larger computing system. Workflow system tool resource usage reporting can help determine the increased resources needed to execute the workflow on larger systems. For researchers without access to free or granted computing resources, this strategy can save significant cost.

#### Gain quick insights using sketching algorithms

Understanding the basic structure of data, the relationship between samples, and the approximate composition of each sample can very helpful at the beginning of data analysis, and can often drive analysis decisions in different directions than those originally intended. Although most bioinformatics workflows generate these types of insights, there are a few tools that do so rapidly, allowing the user to generate quick hypotheses that can be further tested by more extensive, fine-grained analyses. Sketching algorithms work with compressed approximate representations of sequencing data and thereby reduce runtimes and computational resources. These approximate representations retain enough information about the original sequence to recapitulate the main findings from many exact but computationally intensive workflows. Most sketching algorithms estimate sequence similarity in some way, allowing you to gain insights from these comparisons. For example, sketching algorithms can be used to estimate all-by-all sample similarity which can be visualized as a Principal Component Analysis or a multidimensional scaling plot, or can be used to build a phylogenetic tree with accurate topology. Sketching algorithms also dramatically reduce the runtime for comparisons against databases (e.g. all of GenBank), allowing users to quickly compare their data against large public databases.

Rowe 2019 [113] reviewed programs and genomic use cases for sketching algorithms, and provided a series of tutorial workbooks (e.g. Sample QC notebook: [114]).

#### Use the right tools for your question

RNA-seq analysis approaches like differential expression or transcript clustering rely on transcript or gene counts. Many tools can be used to generate these counts by quantifying the number of reads that overlap with each transcript or gene. For example, tools like STAR and HISAT2 produce alignments that can be post-processed to generate per-transcript read counts [115,116]. However, these tools generate information-rich output, specifying per-base alignments for each read. If you are only interested in read quantification, quasi-mapping tools provide the desired results while reducing the time and resources needed to generate and store read count information [117,118].

#### Seek help when you need it

In some cases, you may find that your accessible computing system is ill-equipped to handle the type or scope of your analysis. Depending on the system, staff members may be able to help direct you to properly scale your workflow to available resources, or guide you in tailoring computational unit allocations or purchases to match your needs.

## Strategies for troubleshooting

Workflows, and research software in general, invariably require troubleshooting and iteration. When first starting with a workflow system, it can be difficult to interpret code and usage errors from unfamiliar tools or languages [2]. Further, the iterative development process of research software means functionality may change, new features may be added, or documentation may be out of date [119]. The challenges of learning and interacting with research software require time and patience [4].

One of the largest barriers to surmounting these challenges is learning how, when, and where to ask for help. Below we outline a strategy for troubleshooting that can help build your own knowledge while respecting both your own time and that of research software developers and the larger bioinformatic community. In the “where to seek help” section, we also recommend locations for asking general questions around data-intensive analysis, including discussion of tool choice, parameter selection, and other analysis strategies. Beyond these tips, workshops and materials from training organizations such as the Carpentries, R-Ladies, RStudio can arm you with the tools you need to start troubleshooting and jump-start software and data literacy in your community [120]. Getting involved with these workshops and communities not only provides educational benefits but also networking and career-building opportunities.

### How to help yourself: Try to pinpoint your issue or error

Software errors can be the result of syntax errors, dependency issues, operating system conflicts, bugs in the software, problems with the input data, and many other issues. Running the software on the provided test data can help narrow the scope of error sources: if the test data successfully runs, the command is likely free of syntax errors, the source code is functioning, and the tool is likely interacting appropriately with dependencies and the operating system. If the test data runs but the tool still produces an error when run with your data and parameters, the error message can be helpful in discovering the cause of the error. In many cases, the error you’ve encountered has been encountered many times before, and searching for the error online can turn up a working solution. If there is a software issue tracker for the software (e.g. on the GitHub, GitLab, or Bitbucket repository), or a Gitter, Slack, or Google Groups page, performing a targeted search with the error message may provide additional context or a solution for the error. If targeted searches do not return a results, Googling the error message with the program name is a good next step. Searching with several variants and iteratively adding information such as the type of input data, the name of the coding language or computational platform, or other relevant information, can improve the likelihood that a there will be a match. There are a vast array of online resources for bioinformatic help ranging from question sites such as Stack Overflow and BioStars, to personal or academic blogs and even tutorials and lessons written by experts in the field [121]. This increases the discoverability of error messages and their solutions.

Sometimes, programs fail without outputting an error message. In cases like these, the software’s help (usually accessible on the command line via `tool-name --help`) and official documentation may provide clues or additional example use cases that may be helpful in resolving an error. Syntax errors are extremely common, and typos as small as a single, misplaced character or amount of whitespace can affect the code. If a command matches the documentation and appears syntactically correct, the software version (often accessible at the command line `tool-name --version`) may be causing the error.

Best practices for software development follow “semantic versioning” principles, which aim to keep the arguments and functionality the same for all minor releases of the program (e.g. 1.1 to 1.2) and only change functions with major releases (e.g. 1.x to 2.0).

### How to seek help: include the right details with your question

When searching for the error message and reading the documentation do not resolve an error, it is usually appropriate to for seek help either from the software developers or from a bioinformatics community. When asking for help, it’s essential to provide the right details so that other users and developers can understand the exact conditions that produced the error. At minimum, include the name and version of the program, the method used to install it, whether or not the test data ran, the exact code that produced the error, the error message, and the full output text from the run (if any is produced). The type and version of the operating system you are using is also helpful to include.

Sometimes, this is enough information for others to spot the error. However, if it appears that there may bug in the underlying code, specifying or providing the minimum amount of data required to reproduce the error (e.g. reproducible example [122,123]) enables other to reproduce and potentially solve the error at hand. Putting the effort into gathering this information both increases your own understanding of the problem and makes it easier and faster for others to help solve your issue. Furthermore, it signals respect for the time that these developers and community members dedicate to helping troubleshoot and solve user issues.

### Where to seek help: online and local communities of practice

Online communities and forums are a rich source of archived bioinformatics errors with many helpful community members. For errors with specific programs, often the best place to post is the developers’ preferred location for answering questions and solving errors related to their program. For open source programs on GitHub, GitLab, or Bitbucket, this is often the “Issues” tab within the software repository, but it could alternatively be a Google groups list, gitter page, or other specified forum. Usually, the documentation indicates the best location questions. If question is more general, such as asking about program choice or workflows, forums relevant to your field such as Stack Overflow, BioStars, or SEQanswers are good choices, as posts here are often seen by a large community of researchers. Before posting, search through related topics to double check the question has not already been answered. As more research software development and troubleshooting is happening openly in online repositories, it is becoming more important than ever to follow a code of conduct that promotes open and harassment-free discussion environment [124]. Look for codes of conduct in the online forums you participate in, and make sure you do your part to help ensure a welcoming community for participants of all backgrounds and computational competencies.

While there is lots of help available online, there is no substitute for local communities. Local communities may come in the form of a tech meetup, a users group, a hacky hour, or an informal meetup of researchers using similar tools. While this may seem like just a local version of Stack Overflow, the local, member-only nature can help create a safe and collaborative online space for troubleshooting problems often encountered by your local bioinformatics community. The benefit to beginners is clear: learning the best way to post questions and the important parts of errors, while getting questions answered so they can move forward in their research. Intermediate users may actually find these communities most useful, as they can also accelerate their own troubleshooting skills by helping others solve issues that they have already struggled through. While it can be helpful to have some experts available to help answer questions or to know when to escalate to Stack Overflow or other communities, a collaborative community of practice with members at all experience levels can help all its members move their science forward faster.

If such a community does not yet exist in your area, building this sort of community (discussed in detail in [125]), can be as simple as hosting a seminar series or starting meetup sessions for data analysis co-working. In our experience, it can also be useful to set up a local online forum (e.g. discourse) for group troubleshooting.

## Conclusion

Bioinformatics-focused workflow systems have reshaped data-intensive biology, reducing execution hurdles and empowering biologists to conduct reproducible analyses at the massive scale of data now available. Shared, interoperable research code is enabling biologists to spend less time rewriting common analysis steps, and more time on interesting biological questions. We believe these workflow systems will become increasingly important as dataset size and complexity continue to grow. This manuscript provides a directed set of project, data, and resource management strategies for adopting workflow systems to facilitate and expedite reproducible biological research. While the included data management strategies are tailored to our own experiences in high-throughput sequencing analysis, we hope that these principles enable biologists both within and beyond our field to reap the benefits of workflow-enabled data-intensive biology.

## Acknowledgements

Thank you to all the members and affiliates of the Lab for Data-Intensive Biology at UC Davis for providing valuable feedback on earlier versions of this manuscript and growing these practices alongside us. We also thank the Carpentries Community for fundamentally shaping many of the ideas and practices we cover in this manuscript. This manuscript was collaboratively written using manubot [126] and is available in a GitHub repository [127].

## Author Contributions

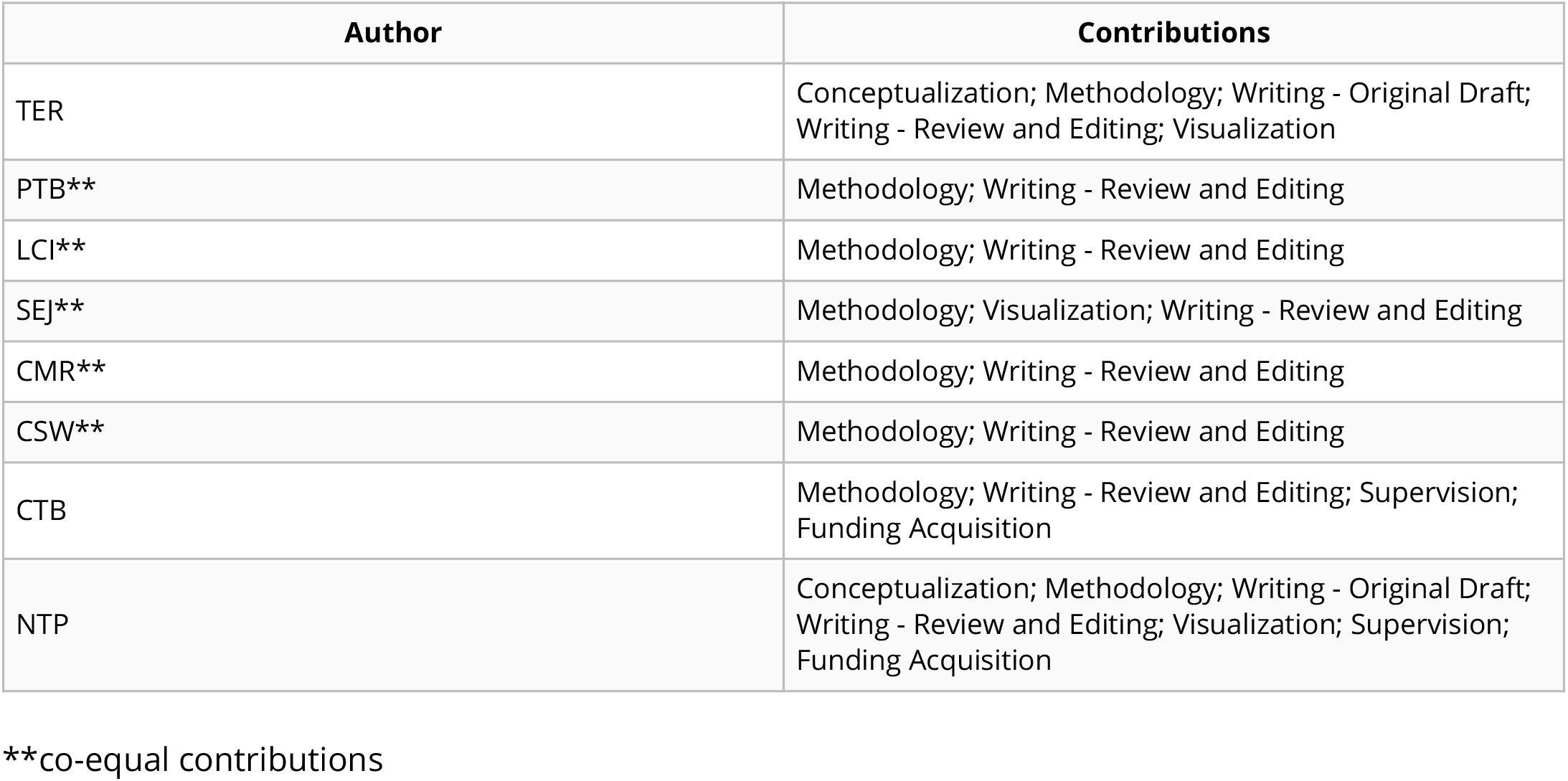

